# Developmental time course of social touch, parvalbumin interneurons, perineuronal nets and Mef2c expression reveals a sensitive period of somatosensory cortex development in prairie voles

**DOI:** 10.1101/2025.07.25.666627

**Authors:** Noah E. P. Milman, Nathan M. McGuire, Jasmine M. Loeung, Lezio S. Bueno-Junior, Carolyn E. Tinsley, Hannah Bronstein, Felice D. Kelly, Peyton T. Wickham, Anjesh Ghimire, Zachary Johnson, Harry Pantazopoulos, Brendon O. Watson, Barbara A. Sorg, Miranda M. Lim

**Author notes:** Correspondence: Miranda M. Lim Department of Neurology, 3181 SW Sam Jackson Park Road, L226 Portland, Oregon 97239-3098. These authors contributed equally (Nathan M. McGuire, Jasmine M. Loeung). Author contributions <u>**CRediT**</u> **Noah E. P. Milman:** conceptualization (equal); funding acquisition (supporting); data curation (lead); formal analysis (lead); investigation (lead); methodology (lead); Software (lead); Visualization (lead); Writing – Original Draft Preparation (lead) **Nathan M. McGuire**: data curation (supporting); formal analysis (supporting); investigation (equal); Writing – Original Draft Preparation (supporting) **Jasmine M. Loeung**: data curation (supporting); formal analysis (supporting); investigation (equal); Writing – Original Draft Preparation (supporting) **Lezio S. Bueno-Junior**: data curation (supporting); formal analysis (supporting); Resources (supporting); Software (supporting); Writing – Original Draft Preparation (supporting) **Carolyn E. Tinsley:** methodology (supporting); Project Administration (supporting); formal analysis (supporting); Resources (supporting); Writing – Original Draft Preparation (supporting) **Hannah Bronstein:** Investigation (supporting); methodology (supporting); Resources (supporting); Software (supporting); Writing – Original Draft Preparation (supporting) **Felice Kelly:** Investigation (supporting); methodology (supporting); Resources (supporting); Software (supporting); Writing – Original Draft Preparation (supporting) **Peyton T. Wickham:** Investigation (supporting); Writing – Original Draft Preparation (supporting) **Anjesh Ghimire:** Software (supporting); Writing – Original Draft Preparation (supporting) **Zachary Johnson:** Supervision (supporting); Writing – Original Draft Preparation (supporting) **Harry Pantazopoulos:** Methodology (supporting); Writing – Original Draft Preparation (supporting) **Brendon O. Watson:** funding acquisition (lead); Supervision (supporting); Writing – Original Draft Preparation (supporting) **Barbara Sorg:** Project Administration (supporting); Supervision (supporting); Writing – Original Draft Preparation (supporting) **Miranda M. Lim:** conceptualization (equal); funding acquisition (lead); Project Administration (lead); Resources (lead); Supervision (lead); Writing – Original Draft Preparation (supporting).

## Abstract

Social touch facilitates our attachment to others, especially early in life, which may be linked to the maturation of parvalbumin interneurons (PVI) in the somatosensory cortex (S1). These neurons respond to social touch, mature in a sensory experience-dependent manner, and influence both somatosensory processing and social behavior in models of Autism Spectrum Disorder. Prairie voles (*Microtus ochrogaster*) are an ideal rodent model for studying these concepts since they engage in a species-typical social touch called “huddling”. This study first showed that over development from juvenile to adult, same-sex siblings huddled less and explored more. Next, we tracked two markers of plasticity indicative of PVI maturation, extracellular perineuronal nets (PNNs) and nuclear transcription factor Myocyte enhancing factor 2C (Mef2c) – across seven developmental timepoints. We found that, while PV expression in S1 was stable by P21, PNNs and Mef2c continued to shift afterwards, indicating a protracted development. Four unique clusters of PVIs converge during development between P14-P21, suggesting a sensitive period of PVI development. Finally, to determine environmental factors affecting these processes, environmental enrichment between P21-P28 led to accelerated PVI maturation. This developmental mapping provides a particularly salient model to investigate the molecular underpinnings of cortical and social development.

## Introduction

Social touch can be a display of affiliation, empathy, or consolation that is conserved across species (Chau et al. 2008; Burkett et al. 2016; Goldstein et al. 2018; Sun et al. 2025). However, social touch changes over development as an individual becomes more socially independent. In concert, the somatosensory system, which receives that touch is refined (Bales et al. 2018; Cascio et al. 2019). To date, it is still unclear how refinement (including at the level of brain oscillations, circuit connectivity or molecular composition) relates to specific features of social behavior. A gap exists in our understanding of the exact sensitive period for the development of social touch. Rodents provide an exceptional system to understand this somatosensory processing (Petersen 2019), as represented by whisker-based sensory information, which relay via the thalamus onto layer 4 (L4) of somatosensory cortex barrel fields (hereafter referred to as S1BF) (Yang et al. 2018; Pineda et al. 2025 Jan 10). In fact, individual cells in S1BF can differentiate social and inanimate object touch (Bobrov, 2014).

A subset of inhibitory interneurons, called parvalbumin-containing interneurons (PVIs), are essential for whisker-based sensory processing in rodents (Cardin et al. 2009). Further supporting their role in thalamocortical tactile processing in rodents, PVIs inhibit nearby neurons (Petersen 2019; Yeganeh et al. 2022), filter sensory information (Yu et al. 2016; Yu et al. 2019), are activated at the onset of social touch (Clemens et al. 2019), and they densely populate L4 of S1BF (Rudy et al. 2011; Hammock and Levitt 2012; Lupori et al. 2023). Typical PVI function arises from both genetic programs and sensory experience-dependent maturation (Le Magueresse and Monyer 2013). At the intersection of these processes resides a transcription factor called Myocyte enhancing factor 2C (Mef2c), which underlies PVI differentiation and postnatal survival (Mayer et al. 2018; Moissidis et al. 2024). Mef2c regulates synaptic plasticity and is highly expressed embryonically in the rodent and human cortex, and its expression decays postnatally (Leifer et al. 1993; Leifer et al. 1997; Ward, Sjulson, et al. 2024). Mef2c is a particularly promising candidate for understanding the pathogenesis of neurodevelopmental disorders, and perhaps also as a therapeutic target (Harrington et al. 2016; Ward, Sjulson, et al. 2024).

In addition to the contribution of Mef2c to PVI development, a second developmental hallmark is the appearance of extracellular matrix structures called perineuronal nets (PNNs). PNNs consist of chondroitin sulfate proteoglycan-rich structures surrounding most cortical PVIs (Sorg et al. 2016; Lupori et al. 2023). Both the fast-spiking properties and the survival of PVIs depend on PNNs, whose formation restricts synaptic plasticity across sensory regions, thus closing their critical periods of development (Miyata et al. 2012; Cabungcal et al. 2013; Fawcett et al. 2019). PNN intensity is negatively correlated with excitatory synapses on PVIs and is positively correlated with inhibitory synapses on the same cells (Sigal et al. 2019). PNNs are also formed in a sensory experience-dependent manner, and experimental manipulations that degrade or reshape PNNs in adolescent or adult rodents (i.e., enzymatic digestion or environmental enrichment) are known to reinstate juvenile-like plasticity and behaviors in these animals (Pizzorusso et al. 2002; Nakamura et al. 2009; Schiff et al. 2023). This suggests that PNNs are potential targets for plasticity-enhancing therapeutic approaches. Among all rodent brain regions, S1BF is where PNNs are generally most dense and intense (Lupori et al. 2023), despite species-dependent variations (Nakamura et al. 2009; Dauth et al. 2016). Interestingly, embryonic removal of Mef2c from fate-mapped interneurons leads to reduced PNN density in adult mice (Ward, Nasrallah, et al. 2024), suggesting that Mef2c and PNNs interact during PVI development. However, such interaction between PNNs and Mef2c across postnatal PVI development has not been characterized.

Prairie voles (*Microtus ochrogaster)* are a socially monogamous and biparental rodent species. Due to their species-typical social behavior, these rodents are a powerful model for understanding the neural circuitry of affiliation (McGraw and Young 2010; Kenkel et al. 2021). In contrast to mice, prairie voles also demonstrate species-typical social touch in the form of huddling (side-by-side contact) in novel environments, including with opposite-sex mates and same-sex siblings (Beery et al. 2018; Vahaba et al. 2022). Notably, however, huddling declines with age among same-sex prairie voles (Kelly et al. 2018), suggesting that both social touch and its neural markers undergo longitudinal changes. It remains unclear how the processing of social touch in the somatosensory cortex coincides with these changes. This study aims to describe the sensitive period for social touch using molecular markers of PVI maturation. Here, prairie voles were used to investigate longitudinal changes in: (1) the behavioral display of social touch; and (2) extracellular (PNN) and nuclear (Mef2c) markers of cortical parvalbumin development. This study is the first to describe the expression of canonical sensitive period molecules in the prairie vole, demonstrating the third postnatal week as a sensitive period of development. Thus, we provide a framework to understand the cellular basis of species-typical social touch in this highly social model organism.

## Materials and Methods

### Subjects

Subjects were male and female prairie voles bred and housed at the Veterans Affairs Portland Health Care System and managed by the Veterinary Medical Unit (VMU) Staff. Breeder pairs within the colony originated from a diverse pool of wild-captured prairie voles and exchanged among several institutions, including the University of Colorado Boulder, Florida State University, and University of California, Davis. Breeder pairs were checked daily at 0700-0800 h for the presence of newborn pups. The day of birth is considered postnatal day 1 (P1). The animals used in this study were sourced from 58 unique litters and 24 breeder pairs, to capture a genetically and behaviorally heterogeneous sample. Colony rooms were set at a 14:10 hours lights on (0700), lights off (2100) cycle at a temperature between 68 – 71 degrees Fahrenheit. Weekly cage changes did not coincide with the day of any experimental recordings described in this paper.

At P21, animals were weaned into appropriate housing conditions in cages of two to three same sex siblings. Housing conditions include: VMU housing (rolled-paper bedding, one-chew toy, two-nestlets) [Experiment 1 behavior], standard housing (SH) (2000 mL corn-cob bedding, one-chew toy, two-nestlets) or enriched environment (EE), (4000 mL of corn cob bedding, chew-toy, plastic tent, plastic igloo, paper-toilet roll, four nestlets and a hanging water bottle) [Experiment 1,2 and 3 immunohistochemistry]. The EE protocol lasted for 7 days while other housing conditions were consistent throughout the study.

### Social behavior test

All animals were naïve to any experimental manipulation present in our colony. To capture behavior, a monochrome FLIR Blackfly S USB3 camera (0.4 MP, Sony IMX287 sensor) was used, with a resolution of 720 X 540 pixels at 30 fps mounted with a 6 mm Edmunds Optics lens. Four files were acquired simultaneously using StreamPix (version 9.0).

All animals were housed in the VMU condition. After a thirty-minute habituation period to the room, siblings were placed in a clean cage containing paper-roll bedding for thirty minutes, and behavior was subsequently characterized using LabGym2 (Figure 1A).

**Figure 1:**
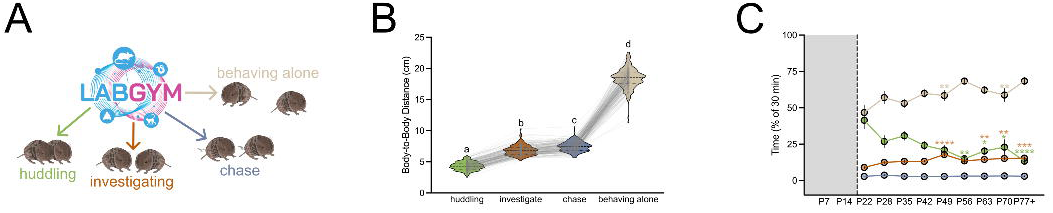
Prairie vole social interactions shift toward independence with increasing age. A. Schematic of experimental design and example LabGym. Epochs were categorized as huddling (side-by-side contact), investigation (including sniffing/being-sniffed, or approaching), chase, or as behaving alone (including idling in place or movement separate from their sibling). B. Difference between animals in mean body-centered coordinates for each behavioral state, lines represent individual sibling dyads. Letters represent significant post-hoc comparisons between behavioral categories. C. Mean proportion of 30 minute recording in each behavioral state (behaving alone, beige; huddling, green; investigate, orange; chase, blue) across postnatal age, data presented as mean +/- standard error of the mean. Asterisks represent significant multiple comparisons vs the P22 timepoint.

### Behavioral annotation

LabGym2-based behavioral annotations followed the ethological criteria listed below, as described previously (Goss et al. 2024).

*Approaching*: A prairie vole is moving toward the conspecific that is idling in place.

*Sniffing*: A prairie vole is sniffing or closely investigating the conspecific.

*Being sniffed*: A prairie vole is being sniffed or closely investigated by the conspecific.

*Huddling*: The two prairie voles are immobile or showing small in-place movements while maintaining side-by-side body contact regardless of body direction.

*Moving*: A prairie vole is moving alone without interacting with the conspecific.

*Idling*: A prairie vole is immobile or showing small in-place movements without interacting with the conspecific.

*Chasing*: A prairie vole is moving toward the conspecific that is also moving in the same direction, forming overlapping trajectories.

*Being chased*: A prairie vole is moving away from the conspecific that is also moving in the same direction, forming overlapping trajectories.

Next, in MATLAB (Mathworks), files were trimmed and binned into 1 second epochs and the most probable behavior was determined the “winner” of that epoch. Each epoch was assigned one of four categories: 1) huddling, 2) behaving alone (moving and idling), 3) chase (chasing and being chased) or 4) investigating (approaching, sniffing, and being sniffed). To calculate the cumulative time spent in each behavior, the sum of epochs assigned to each behavior was determined. The average probability of a given behavior across time was highly correlated to the cumulative time spent in the behavior (data not shown, R^2^ > 0.94). Cumulative time spent in each behavior did not pass tests for normality and thus were analyzed with non-parametric one-way ANOVA Kruskal-Wallis test (see Statistics section).

### Tissue collection and Immunohistochemistry

P11 and P14 pups were removed from parents, placed into a new cage, and immediately deeply anesthetized with 5% inhaled isoflurane followed by decapitation and brain harvesting. For all other timepoints beginning at P21 (weaning): after a 30-60 min habituation period following cage change into single housing, animals were deeply anesthetized with 5% isoflurane until loss of consciousness, and brain tissue was harvested. Tissue was placed into ice cold 4% paraformaldehyde (PFA) for 24 hours before transfer into 0.1% sodium azide in 1X Phosphate Buffer Saline (PBS). Sectioning was conducted with a Leica slicing vibratome with coronal sections cut at 30 μm and stored in 0.1% sodium azide in 1X PBS until immunohistochemistry.

Free-floating sections were triple labeled for *Wisteria floribunda* agglutinin (WFA), which recognizes GalNAc residues of the chondroitin sulfate proteoglycans (CSPGs), PV, and Mef2c using immunohistochemistry. Sections were drop fixed in 4% PFA, rinsed with 0.1 M phosphate buffer (PB), and rinsed again in 0.02% Triton x in 1X PBS (0.02% PBS-Tx) three times before 30 min incubation at 80°C in 10% citric acid buffer solution. Next, tissue was rinsed in 0.02% PBS-Tx and incubated in 0.3% H_2_O_2_ in 0.02% PBS-Tx for 30 minutes. Sections were rinsed in 0.02% PBS-Tx three times and incubated in blocking buffer, which was 2% normal goat serum (NGS) in 0.02% PBS-Tx for thirty minutes. Sections were rinsed in 0.02% PBS-Tx three times before incubation with primary antibodies diluted in 2% NGS buffer in 0.02% PBS-Tx for 48 hours in the dark at 4°C 1:1000 biotinylated WFA (Vector labs, #B-1355), 1:4000 mouse anti-PV (Sigma P3088) and 1:500 rabbit anti-Mef2c (ProteinTech 1000-56-1). On the third day, sections were rinsed with 0.02% PBS-Tx three times before a 3 hour room-temperature incubation in the dark with secondary antibodies diluted in 2% NGS: 1:200 Cy2-Streptavidin (Jackson Immuno 016-220-084), 1:300 Cy3 donkey anti-mouse (Jackson Immuno 715-165-150), 1:300 AlexaFluor647 donkey anti-rabbit (Jackson Immuno 715-605-152). Sections were finally washed three times in 0.1 M PB solution and mounted on VWR Frosted Slides with ProLongGold antifade containing DAPI (P36930). Slides were stored at 4°C in light-protected slide-box until imaging.

### Fluorescent slide-scanning (ALMC)

Slides containing tissue at various ages were scanned using identical acquisition parameters on the Zeiss Axioscan7 located at the Advanced Light Microscopy Core (ALMC) at OHSU. Images were captured using a Colibri7 LED light source, Plan-Apo10x 0.45 NA objective, and an Orca Flash4.0 C13440 camera (Hamamatsu). Excitation/Emission filters were as follows; Dapi EX = 390/40, EM = 450/40; AF488 Ex = 470/40, Em = 525/50; AF555 Ex = 550/25, Em = 605/70; AF647 Ex = 640/30 Em = 690/50. Tiled Z-stacks (20 μm range, 4 μm z-interval) were acquired to capture the full tissue with a voxel size (0.7 μm, 0.7 μm, 4 μm). The images were flattened using a maximum intensity extended depth of focus using the ZEN Blue 3.7 software. of 5 slices taken 4 μm apart. These images generated a czi filetype (see image processing below).

### PVI identification and PNN classification

Images were background subtracted across 488, 555 and 647 channels using a filter width of 3320 µm. Regions of interest, including somatosensory cortex and layer 4 of somatosensory cortex were drawn using the pencil tool in Imaris (Oxford, Version 10.1. Using the Surfaces object function in Imaris, PVI soma were detected using 1.3um surface detail smoothing, with background subtraction (local contrast) of 18um. Next, the threshold for background subtraction was set manually according to individual images, using 18.0 µm seed point diameter. Finally, detected PVI surfaces were filtered according to the quality measurement (the brightness of the center of the surface) and number of voxels between 100 and 1000 to remove unlikely surfaces. This batch protocol was run across all images within this dataset.

To determine whether a detected PVI surface was surrounded by a PNN, the distribution of the mean intensity of 488 Channel (WFA) within all PVI surfaces was plotted (see Supplemental Figure 1). Across all detected cells in the dataset, the 25^th^ percentile of WFA signal was 902 AU. The accuracy of this threshold was then verified across images from all age groups. Cells with mean intensity greater than or equal to 902 AU, were deemed PNN+, and those below 902 AU were deemed PNN-.

### Data analysis

Custom R-scripts (RStudio, Version 12.1) were written to process the exported files from Imaris, including:

1. Assignment of PVIs according to hand-drawn region of interest (i.e. Layers 4 “L4” or Layers 2/3. 5, and 6 “non-L4”)
2. PVI classification by PNN+ or PNN-according to 902 AU threshold described above.
3. Merging data from adjacent brain sections from the same animals (technical replicates) to generate averaged results
4. Clustering of PVIs into k clusters (groups) using the kmeans() function. To determine the optimal number of clusters k, we used the gap statistic to compare the total within cluster sum of squares (Supplemental Figure 2A) (Tibshirani et al. 2001).

### Statistics

All assumptions were met for parametric testing, except where described: including behavioral data (Figure 1) which failed assumptions, and therefore we used non-parametric testing (Friedman and Kruskal-Wallis tests). The mean intensity comparisons of Mef2c also failed assumptions for parametric testing and were subsequently log transformed, which allowed for a normal distribution. Otherwise, we utilized two-way ANOVAs with Sidak corrections for multiple comparisons, with a threshold of alpha < 0.05 to reject the null hypothesis. Statistical comparisons were made in GraphPad Prism 10.1.0.

## Results

### Experiment 1: Social behavior across development in same-sex sibling dyads

To examine age related changes in social behavior in prairie voles, same-sex sibling dyads were placed into a novel cage at various timepoints after weaning (P21+).

A social behavior tracker called LabGym (Hu et al. 2023; Goss et al. 2024) was used in conjunction with custom code to label four behaviors: huddling, investigating, chasing, and behaving alone (Figure 1A). For initial verification of LabGym-based annotation, these behaviors were shown to differ in terms of distance between animals [repeated-measures Friedman test: Friedman statistic (4,164) = 464.0, Dunn’s multiple comparison test, each p < 0.0001] (Figure 1B). Then, the incidence of these behaviors was tracked over time, showing that animals spent approximately half of all recorded time behaving alone, which further increased with age [Kruskal-Wallis test for each behavior type; behaving alone: H(9,164) = 23.28 p = 0.003]. The time spent investigating also increased with age [H(9,164)=32.89, p<0.0001], but time spent chasing did not [H(9,164)=4.979, p=0.76] (Figure 1C). Moreover, juveniles (P22) spent more time huddling than did adults (>P49) [H(9,164) = 31.18, p = 0.0001, all post-hoc comparisons p<0.05] (Figure 1C).

Taken together, these data suggest that the time spent engaged in same-sex huddling behavior decreases with postnatal age while time spent independently increases.

### Experiment 2: Changes in parvalbumin interneurons in the somatosensory cortex across development

Based on these age-related changes in social behavior, age-related changes were also investigated in the molecular and cellular systems implicated in social touch. First, in a separate cohort of animals, the typical development of somatosensory cortical PVIs was characterized. PVIs were stained in coronal sections from naïve prairie voles at seven postnatal timepoints [P11, P14, P21 (weaning), P28, P35, P60 and P100+] (Figure 2A). L4 of S1BF was manually identified and measured for density and immunofluorescence (intensity) of PVI somas in L4 versus non-L4 (layers 2/3, 5 and 6), based on the known laminar density of interneurons in S1 (Rudy and Fishell 2010).

**Figure 2:**
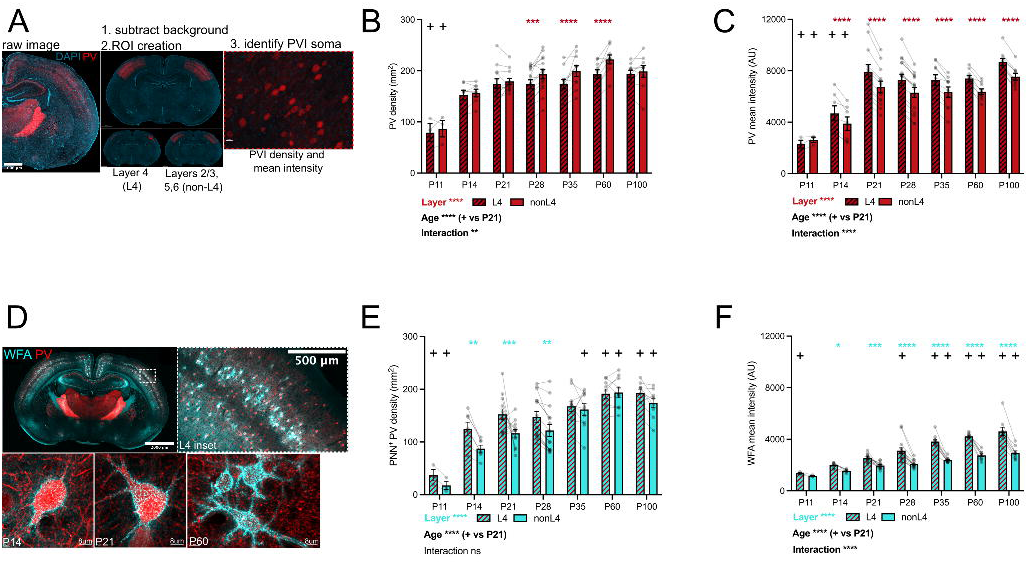
Parvalbumin interneuron and Perineuronal net density intensity in somatosensory cortex of the prairie vole across development. A. Experimental pipeline, left to right: Brain sections imaged on slide scanner, DAPI (blue), PV (red), then background subtracted, followed by hand-drawn ROIs, generating diffuse intensity values in each region (layer 4 “L4” or Layers 2/3, 5, 6 “non-L4”). Inset of L4 PV staining, demonstrating detection of individual PVIs, outlined in white. B. PVI density increases in both L4 (shaded red) and non-L4 (solid red) regions with age. Data presented as mean +/- standard error of the mean, circles represent individual animals. + symbol represent significantly different value compared to P21 weaning timepoint. Asterisks represent significant difference between layers (p<0.05). C. PV mean intensity within soma of PVIs increases with age and is higher in L4 (shaded red) than non-L4 (solid red) PVIs. Data presented as mean +/- standard error of the mean, circles represent individual animals. + symbol represent significantly different value compared to P21 weaning timepoint. Asterisks represent significant difference between layers (p<0.05). D. Top left: PNN expression in the developing prairie vole, WFA (cyan), PV (red), 2mm scale bar. Top right, inset of somatosensory cortex, scale bar 500μm. Bottom: Representative confocal 60x images of PVIs at P14, P21 and P60, demonstrating age-dependent increase in the coverage of the PVI. Scale bar, bottom left, 8μm. E. Density of PNN+ PVIs by age in L4 (shaded cyan) and non-L4 (solid cyan) increases with age. Data presented as mean +/- standard error of the mean, circles represent individual animals.+ symbol represent significantly different value compared to P21 weaning timepoint (p<0.05). F. WFA mean intensity increases with age and is higher in L4 (shaded cyan) than non-L4 (solid cyan). Data presented as mean +/- standard error of the mean, circles represent individual animals. + symbol represent significantly different value compared to P21 weaning timepoint (p<0.05).

PVI density was found to plateau by P14, with greater density in non-L4 than L4 at P28, P35 and P60 [repeated-measures 2-way ANOVA, within subjects effect of layer F(1,58) = 29.14, p < 0.0001, between subjects effect of postnatal age, F(6,58) = 9.021, p < 0.0001, and interaction, F(6,58) = 3.201, p = 0.0088, with Sidak multiple comparisons] (Figure 2B). PV protein expression, represented by PV mean intensity within PVIs, was found to plateau by P21, with L4 showing higher PV expression compared to non-L4 at all timepoints except P11 [repeated-measures 2-way ANOVA, within subject effect of layer F(1,58) = 262.6, p < 0.0001, between subject effect of postnatal age, F(6,58) = 10.52, p < 0.0001 and interaction F(6,58) = 7.712, p < 0.0001, with Sidak multiple comparisons] (Figure 2C). In summary, PVIs in prairie vole S1BF were present by P14, but PV protein expression was highest in L4, suggesting a unique activity profile of these cells.

The PNN sheath surrounding PVIs restrict synapse formation, with the intensity of PNN labeling correlating negatively with the size of physical locations for synapses to form (Sigal et al. 2019). Here, PNNs in S1BF were labeled with WFA (Figure 2D). PVIs positive for PNN (PNN+) and negative for PNN (PNN-) were compared based on the mean intensity surrounding the cell soma (Supplemental Figure 1, Figure 2A). In contrast to PVI density, which was stable by P14, there was an increase in PNN+ PVI density in adulthood (P60 and P100) [repeated-measures 2-way ANOVA, within subject effect of layer, F(1,58) =21.88, p < 0.0001, between subject effect of postnatal age F(6,58) = 18.85, p < 0.0001 and a trend for an interaction F(6,58) = 2.108, p = 0.066, with Sidak multiple comparisons] (Figure 2E). In addition, PNNs were not uniformly distributed, as L4 showed a greater density of PNN+ PVIs than non-L4 at P14, P21, and P28, (Figure 2E). Among PNN+ PVIs, WFA mean intensity was greater in L4 than non-L4 at all timepoints, except P11 [repeated-measures 2-way ANOVA, within subject effect of layer, F(1,58) = 189.2, p < 0.0001, between subject effect of postnatal age F(6,58) = 31.51, p < 0.0001 and interaction F(6,58) = 8.901, p < 0.0001, with Sidak multiple comparisons] (Figure 2F). Taken together, these results suggest that PVI maturation, as indicated by increased density and intensity of PNNs surrounding PVI soma, occurs in a protracted manner. PNNs appear first in L4 followed by non-L4, and the extended period during which PNNs condense may enable synaptic restructuring into adulthood.

Activity-dependent genetic programs support PVI maturation, including nuclear expression of transcription factor Mef2c (Moissidis et al. 2024; Ward, Nasrallah, et al. 2024). Here, high levels of Mef2c expression were observed in the developing prairie vole somatosensory cortex, including its co-localization in PVIs (Figure 3A-B). Mef2c expression within PVIs decreased with postnatal age and was greater in L4 compared to non-L4 at all timepoints [repeated-measures 2-way ANOVA, within subject effect of layer, F(1,58) = 396.9, p < 0.0001, between subject effect of postnatal age, F(6,58) = 8.1, p < 0.0001 and interaction F(6,58) = 12.88, p < 0.0001] (Figure 3C). To understand the relationship between extracellular PNNs and nuclear Mef2c, PNN+ PVIs (teal) and PNN-PVIs (red) were compared. Few PNN-PVIs were identified in adult prairie voles, with PNN+ PVIs being more numerous than PNN-PVIs by P14 [repeated measures 2-way ANOVA, within subject effect of PNN class F(1,58) = 237.8, p < 0.0001, between subject effect of postnatal age F (6,58) = 9.353, p < 0.0001, and interaction, F(6,58) = 18.43, p < 0.0001, with Sidak multiple comparisons] (Figure 3D). By comparing PNN+ and PNN-PVIs, higher levels of Mef2c among PNN-PVIs were found at all postnatal ages except P11, when few PVIs show a fully-formed PNN [repeated-measures 2-way ANOVA, within subject effect of PNN class, F(1,58) = 1024, p < 0.0001, between subject effect of postnatal age, F(6,58) =4.685, p = 0.0006, and interaction, F(6,58) = 6.572, p<0.0001, with Sidak multiple comparisons] (Figure 3E). Consistent with this, lower PV mean intensity in PNN-PVIs was observed at all ages. Furthermore, this effect was found to strengthen with age [repeated measures 2-way ANOVA, within subject effect of PNN class, F(1,58) = 411.0, p < 0.0001, between subject effect of postnatal age F(6,58) = 9.192, p< 0.0001, but no significant interaction, F(6,58) = 1.557, p = 0.1761, with Sidak’s multiple comparisons] (Figure 3F).

**Figure 3:**
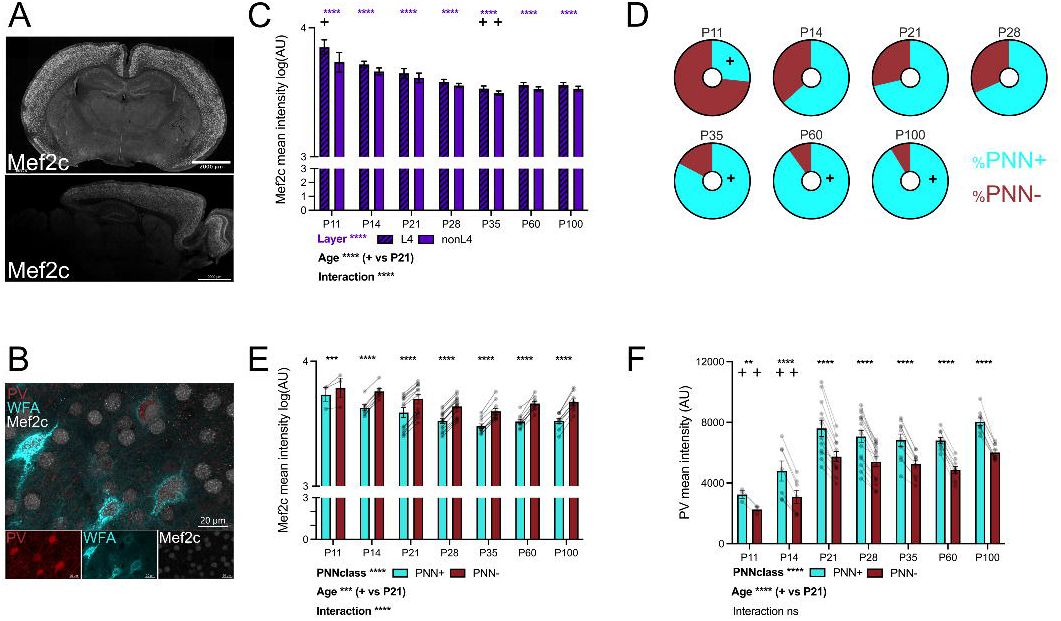
Immature PVIs in S1BF have high nuclear Mef2c, low PV and low PNN expression. A. Mef2c expression in white (coronal, top, and sagittal, bottom), scale bar is 2mm. B. Representative 60x confocal image demonstrating nuclear colocalization of Mef2c (white) and PV (red), and WFA (cyan), channels split below. Scale bar, 20μm. C. Mef2c mean intensity within PVIs decreases with age and is significantly increased in L4 compared to non-L4. Data presented as mean +/- standard error of the mean. + symbol represent significantly different value compared to P21 weaning timepoint (p<0.05). D. The proportion of PVIs in S1BF without a PNN (PNN-, red), declines with age while PVIs with a PNN (PNN+, cyan) increase. Data presented as mean proportion of PVIs classified as PNN+ and PNN-by postnatal age. + symbol represent significantly different value compared to P21 weaning timepoint (p<0.05). E. Mef2c nuclear expression is increased in PVIs lacking a PNN (PNN-, red), at all age timepoints, except P11. Data presented as mean +/- standard error of the mean, circles represent individual animals. + symbol represent significantly different value compared to P21 weaning timepoint (p<0.05). F. PV mean intensity is decreased in PVIs lacking a PNN (PNN-, red) at all age timepoints. Data presented as mean +/- standard error of the mean, circles represent individual animals. + symbol represent significantly different value compared to P21 weaning timepoint (p<0.05).

To decipher the precise timing of PVI maturation, we next grouped individual PVIs from our dataset in an unsupervised manner using kmeans clustering based on PV and Mef2c mean expression. Clustering in this manner allowed us to determine variance in molecular composition regardless of the layer of somatosensory cortex, age or identity of each animal. We found four unique populations of PVIs described by low to medium PV and high Mef2c “Cluster 1” (grey), low PV and low Mef2c “Cluster 2” (green), medium PV and low Mef2c “Cluster 3” (purple) and high PV and low Mef2c “Cluster 4” (orange) within the entire dataset, N = 208,208 PVIs from 75 animals (Figure 4A). The proportion of PVIs in clusters 1 and 2 declined with age, while cluster 3 demonstrated a linear increase with age (Figure 4B, Supplemental Figure 2B) [repeated measures 2-way ANOVA, significant within subject effect of Cluster id, F(1.618,93.84)=20.11, p<0.0001, no significant between subject effect of postnatal age, F(6,58)=1.096, p=0.3759, and a significant interaction, F(18,174)=5.759, p<0.0001, with Sidak multiple comparisons] (Figure 4B). P21 represented the inflection point whereby clusters 2 and 3 crossed and the majority of PVIs belonged to cluster 3. PVIs within these clusters could also be distinguished based on the mean intensity of PNNs surrounding them, with increasing WFA mean intensity in each cluster as well as increased proportion of PVIs surrounded by a PNN [repeated measures 2-way ANOVA, significant within subject effect of cluster id, F(2.129,123.5)=395.3, p<0.0001, significant between subject effect of postnatal age, F(6,58)=18.82, p<0.0001, and a significant interaction, F(18,174)=7.480, p<0.0001, with Sidak multiple comparisons] (Figure 4C, Supplemental Figure 2C). Together these findings suggest that PVI maturation as described by differences in mean intensity of PV, nuclear Mef2c and extracellular WFA shift in composition at P21.

**Figure 4:**
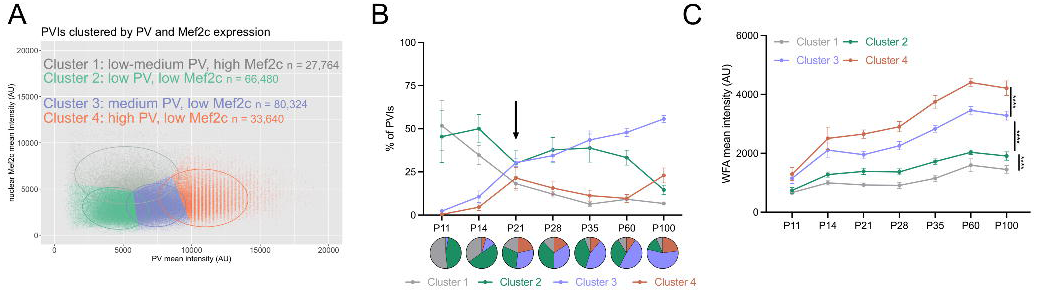
Four unique clusters of PVIs converge during development between P14-P21, suggesting a sensitive period of PVI development. A. Individual PVIs clustered by PV (x-axis) and nuclear Mef2c (y-axis) expression, total N = 208,208 PVIs, from 75 animals. K-means cluster generated 4 clusters Cluster 1 (gray): low-medium PV, high Mef2c (N = 27,764), Cluster 2 (green): low PV, low Mef2c (N=66,480), Cluster 3 (purple): medium PV, low Mef2c (N=80,324), and Cluster 4 orange): high PV, low Mef2c (N=33,640). B. Proportion of PVIs belonging to each cluster by postnatal age. Data presented as mean +/- standard error of the mean, with pie chart below each age. C. WFA mean intensity according to cluster and postnatal age. Data presented as mean +/- standard error of the mean. Asterisks represent significant post-hoc comparisons of cluster id (p<0.0001).

### Experiment 3: Post-weaning enriched environment

Studies in traditional laboratory rodents have shown that enriched environments (EE) modulate PNN formation in adolescents and reshape PNN expression in adults or studies of neurodegeneration. Thus, the timing and duration of enrichment is essential for studying the contribution of EE on S1 development (Bartoletti et al. 2004; Nithianantharajah and Hannan 2006; Greifzu et al. 2014; Slaker et al. 2016a). Experiment 3 examines this phenomenon in prairie voles.

Prairie voles were subjected to EE in the home cage one week after weaning (P21-P28). Then, PNNs or markers of PVI maturity were examined in EE compared to standard housed animals (Figure 5A). EE was found to significantly increase the number of PNN+ PVIs in both L4 and non-L4 regions [repeated-measures 2-way ANOVA, within subject effect of layer (F(1,23)=14.12, p=0.001, between subject effect of housing (F(1,23)=7.665, p=0.0109), and no significant interaction F(1,23)=0.1645, p = 0.6888, with Sidak multiple comparisons) (Figure 5B). The mean intensity of PV was increased after EE, with a significant increase in L4 compared to non-L4 [repeated-measures 2-way ANOVA, within subject effect of layer, F(1,23)=202.5, p<0.0001, between subject effect of housing, F(1,23) = 5.265, p=0.0312) and interaction F(1,23)=5.068, p=0.0342, with Sidak’s multiple comparisons] (Figure 5C). In addition, EE slightly the mean intensity of PNNs surrounding PVIs in L4 but not in non-L4 [repeated-measures 2-way ANOVA, within subject effect of layer, F(1,23)=133.3, p<0.0001, and a trend for between subject effect of housing, F(1,23)=4.201, p=0.052, but no interaction, F(1,23) =1.491, p=0.2344, with Sidak multiple comparisons] (Figure 5D). Nuclear Mef2c expression was not changed after EE but was significantly higher in L4 than non-L4 regions, thus replicating earlier results [repeated-measures 2-way ANOVA, within subject effect of layer, F(1,23)=129.9, p<0.0001, no between subject effect of housing F(1,23)=0.01089, p=0.9178, or interaction, F(1,23)=2.561, p=0.1232] (Figure 5E). One week of enriched environment reduced the proportion of PVIs in cluster 2, and increased those in clusters 3 and 4 [repeated measures 2-way ANOVA, significant within subject effect of Cluster id, F(3,69)=10.6, p<0.0001, no significant between subject effect of housing, F(1,23)=0.02521, p=0.8752, and a significant interaction, F(3,69)=3.353, p=0.0238, with Sidak multiple comparisons] (Figure 5F).

**Figure 5:**
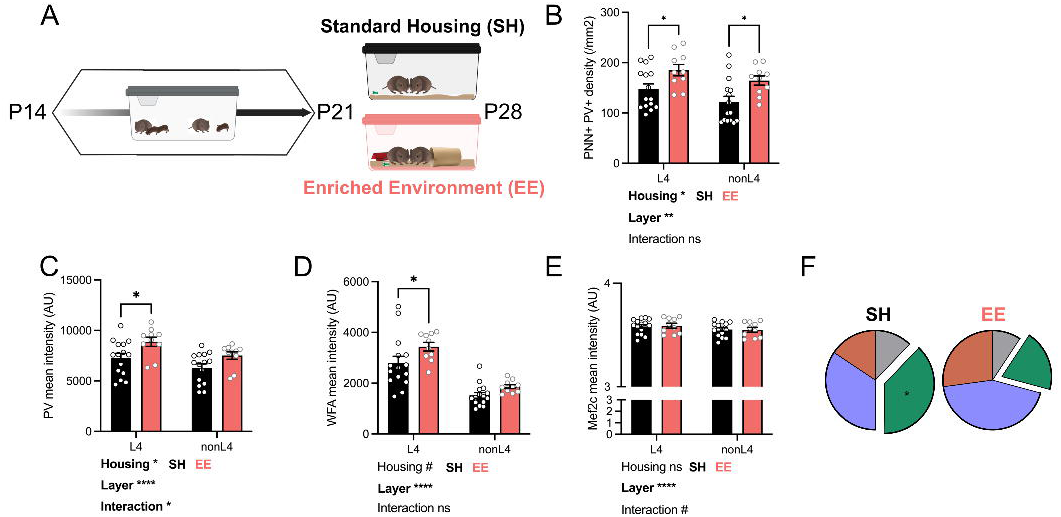
EE accelerates formation of PNNs surrounding PVIs and PVI maturity. A. Schematic of experimental protocol: animals are weaned at P21 into standard housing (SH, black) or environmental enrichment (EE, peach), and brain tissue is harvested one week later at P28, see Methods. B. The density of PNN+ PVIs is significantly increased by EE in both L4 and non-L4 regions. Data presented as mean +/- standard error of the mean, circles represent individual animals. Asterisks represent significant post-hoc comparisons of housing group (p<0.05). C. PV mean intensity is increased by EE in L4, but not in non-L4. Data presented as mean +/- standard error of the mean, circles represent individual animals. Asterisks represent significant post-hoc comparisons of housing group (p<0.05). D. WFA mean intensity surrounding PVIs is increased by EE in L4, but not non-L4. E. Nuclear Mef2c mean intensity is no different between EE and SH in either region, but expression is higher in L4. Data presented as mean +/- standard error of the mean, circles represent individual animals. F. Proportion of PVIs belonging to each cluster according to housing group. Data presented as mean proportion of each cluster by group. Asterisks represent significant post-hoc comparisons of housing in cluster 2 (p<0.05).

Taken together, these data demonstrate that even a short period of sensory enrichment can accelerate PVI maturation in a layer-specific manner in prairie voles.

## Discussion

Two developmental biology dimensions were examined here, one behavioral and another cellular, both in developing prairie voles. First, the typical development of social interactions among prairie vole siblings was quantified using computational tracking, showing a decay in social touch and a proportional increase in non-social exploration as the animals grew older. Then, this shift in social behavior was found to occur following parvalbumin interneuron maturation in the somatosensory cortex. In this maturation process, PVIs were observed to mature first in L4 and then non-L4 cortical layers, suggesting orderly neuroanatomical development. The most substantial shift in PVI maturation occurred in the third postnatal week, but PNNs and Mef2c continued to shift into adulthood, supporting a sensitive period of development. Our findings also indicate heterogeneity within the population of PVIs, with a subset of these cells maintaining plasticity into adulthood, as biomarked by low extracellular PNN and high nuclear Mef2c expression. Finally, environmental enrichment was found to accelerate the maturation of PVIs, particularly in L4 of the somatosensory cortex, indicating that the orderly maturation mentioned above is influenced by sensory experience. These results span key developmental windows (juvenile, adolescence, early-adulthood and adulthood) and are neuroanatomically specific, laying groundwork for studies on somatosensory development, including in studies relevant to Autism Spectrum Disorder (ASD).

### Change in social touch reflects a shift toward independence

Social touch from both parents and siblings are essential in altricial species, including humans, as social touch is believed to scaffold the development of sensory processing (Maitre et al. 2017; Bales et al. 2018; Krol et al. 2019). However, in humans, the nature, intention, and meaning of social touch may evolve as a child becomes more independent (Cascio et al. 2019). Rodents demonstrate an age-dependent shift in social behavior, first in their capacity to sense familiar conspecifics and later in early-adulthood, in their preference to interact (Naskar et al. 2019; Nardou et al. 2023; Florbela de Rocha-Almeida). In parallel, in the present study, a reduction in huddling and an increase in solitary behaviors was observed in young adult prairie voles, and in this species, the shift in independence occurs near the time of sexual maturity, typically around P45 in males (Mateo et al. 1994) and P30-45 in females (Solomon 1991).

Interestingly, this timing coincides with the typical period of natal dispersal observed in the wild (McGuire et al. 1993). Previously, Kelly and colleagues reported a decline in huddling among novel male-male prairie vole dyads from pre-weaning to post-weaning timepoints (Kelly et al. 2018). This is consistent with what we would hypothesize based on PVI maturation between those timepoints. Together, the present results provide context to understand how developmental exposures, including the loss of sleep during a sensitive window of brain development (P14-P21) in prairie voles may impact the expression of both peer (same-sex sibling) or pair (opposite-sex partner) bonding (Jones et al. 2019; Bueno-Junior et al. 2023).

### A sensitive period for PVI maturation in the somatosensory cortex of prairie voles

Changes in PVI density, PV expression and function are a potential mechanism contributing to ASD during development (Filice et al. 2016; Contractor et al. 2021; Wood et al. 2021). To better understand how cortical interneurons that support sensory processing and social touch mature alongside these changes in huddling, we assessed PVI maturation in the somatosensory cortex of prairie voles. The population of PVIs reached adult-levels by P14 already, but PVIs continued to mature with respect to the expression of PV, extracellular PNNs and nuclear Mef2c into later ages. Our further analysis of more than 200,000 unique PVIs suggests four developmentally regulated clusters of PVIs according to features of PV mean intensity and Mef2c expression, the first two representing “plastic” and the final two representing “fixed” states. Notably, PV expression increased until P21, as did membership in more “fixed” clusters 3 and 4, implying a sensitive period of PVI development. These findings in prairie voles are consistent with the approximate developmental timeline in mice, in which PVIs appear histologically and exhibit mature firing properties by P21 (Nowicka et al. 2009; Pouchelon et al. 2021; Kon et al. 2024).

The present study provides context to a sensitive period of PVI maturation, as early-life sleep disruption between P14-P21 increases the density of PVIs in L4 of S1BF of adult prairie voles (Jones et al. 2019). Together, the maturation of PVIs is implicated in the maintenance of the excitatory:inhibitory (E:I) ratio in the brain, which may be the mechanistic link between loss of sleep and behavioral features of ASD (Bridi and Peixoto 2025).

Postnatal experience can impact the activity and formation of PVIs and the synaptic connections made within the circuit that support complex sensory processing and social touch. Expectedly, PNNs, which restrict synapses, mature in an age-dependent manner in prairie voles. In this study, the number of PVIs containing a PNN (PNN+ PVIs) increased first in L4 then non-L4 regions, and the intensity of PNNs increased in L4 relative to non-L4. This is consistent with studies in mice showing a high density of PNNs in L4 of S1BF along with age-dependent increases in mean intensity of PNNs (Nowicka et al. 2009; Ueno et al. 2019; Lupori et al. 2023). We also observed increases in PNN density and mean intensity in early-adulthood. However, because anatomical features, including the appearance of PNNs in the somatosensory cortex occurs earlier than in other sensory circuits, it has been proposed that it is the first critical period to close (before the third postnatal week) (Erzurumlu and Gaspar 2012; Lo et al. 2017; Reh et al. 2020; Ramsaran et al. 2025). Conversely, protracted development of PVIs and regional heterogeneity in PNN expression could enable plasticity into adulthood and dynamic PVI functions, which are not appropriately described by a critical period (Koelbl et al. 2015; Dauth et al. 2016; Lau et al. 2020). In our study, we find PVIs in the somatosensory cortex of the prairie vole undergo protracted development, with heightened sensitivity during the third postnatal week, supporting a sensitive period instead.

Mef2c has been implicated in the neuropathogenesis of ASD, potentially through its role in synaptic refinement (Barbosa et al. 2008; Putman et al. 2024) and influence on PVI functional maturation (Harrington et al. 2016; Mayer et al. 2018; Moissidis et al. 2024; Ward, Nasrallah, et al. 2024). Interestingly, PVIs identified here as lacking a PNN showed higher expression levels of Mef2c into adulthood, when Mef2c is typically downregulated in the cortex. Consistent with the idea that PNNs and PVIs mature together, our study shows that PVIs lacking a PNN had lower expression of PV compared to neighboring neurons with a PNN. Moreover, PVIs belonging to clusters 1 and 2 had significantly lower mean expression of PNNs. This unbiased profiling of PVIs implies that each PVI may have a unique molecular makeup, as suggested by Devienne and colleagues (Devienne et al. 2021; Wang et al. 2023). Future experimental manipulations to alter this highly plastic population of PVIs could mechanistically test the function of these cells in adults. Defining PVI maturation based on multiple features, including Mef2c expression, sharpens our capacity to identify sensitive periods of brain development.

### Factors that influence PVI maturation

Identifying factors that drive, delay or accelerate sensory development is of crucial importance to designing translationally-relevant and accessible interventions for individuals with differences in sensory processing and social interactions. We interrogated whether EE from P21 to P28 (1 week post-weaning) could accelerate the maturation of PVIs in prairie voles. EE increased the density of PNN+ PVIs in both L4 and non-L4 regions. Meanwhile, EE increased the mean intensity of both PV and PNNs in L4 compared to standard housed prairie voles. These findings suggest that EE has the capacity to reshape the typical composition of the somatosensory cortex at P28. PVIs require adequate sensory experience to mature (Nowicka et al. 2009; Yang et al. 2018), and enriched environments can reshape adult PVIs (Madinier et al. 2014; Slaker et al. 2016b; Huang et al. 2021). EE is a promising non-invasive intervention for individuals with ASD who process tactile information differently, especially in the context of social touch (Mammen et al. 2015; Kaiser et al. 2016). These findings have been expanded on in mouse models of ASD, demonstrating aberrant sensory processing and avoidance to both tactile and social stimuli, which may be remedied with a GABA-A receptor agonist (Orefice et al. 2019; Chari et al. 2023). EE in rodent studies can identify mechanisms underlying the benefits of sensory-integration therapy in ASD on improving social behavior (Reynolds et al. 2010; Aronoff et al. 2016).

One limitation of this study is that not all timepoints across development were sufficiently powered to investigate sex as a biological factor, and the sexes were represented equally and pooled together. It would be useful to characterize any differences in the timing of PVI maturation as it relates to the onset of differences in sensory processing and social behavior between females and males. PVI activity is regulated by the estrus cycle and estrogen, highlighting that PVIs may be particularly sensitive to sex-specific hormonal signaling, e.g. during adolescence (Wu et al. 2014; Clemens et al. 2019). Similarly, PNN expression is regulated by the estrus cycle (Laham et al. 2022). Ultimately, it would also be useful to manipulate the activity of PVIs and PNNs *in vivo* and investigate the consequences for social touch in prairie voles.

### Concluding remarks

Our study supports a sensitive period of PVI maturation that can be modulated by enriched sensory experience in the prairie vole. Coinciding with this cortical maturation process, the incidence of huddling among siblings declined with age. Using these longitudinal development “templates”, future investigations of early-life experiences that modify somatosensory cortex maturation are needed to better understand atypical social development, thus identifying targets for translationally-relevant interventions.

## Supporting information

Supplemental Materials including figures

## Acknowledgements

N.E.P.M. thanks the generous support of Achievement Reward for College Scientists (ARCS) Oregon Chapter. We appreciate the technical and imaging support from Dr. Brian Jenkins and Dr. Stefanie Kaech Petrie at the OHSU Advanced Light Microscopy Core Imaging Core. The contents do not represent the views of the U.S. Department of VA or the United States Government. Please send all correspondence to Dr. Miranda Lim

## Conflict of interest

The authors declare no competing financial interests.

## Funding sources

This research was supported by NIH grants F31MH136684 and T32AG055378 (N.E.P.M.), MH131592 (B.O.W., M.M.L., and L.S.B-J), R35GM155357 (ZVJ) and OD P51OD011132 to Emory National Primate Center, DA055645, NS131645, and Good Samaritan Foundation of Legacy Health (B.A.S.), and Portland VA Research Foundation (M.M.L).

## Notes

### Competing Interest Statement

The authors have declared no competing interest.

