## Supplemental Materials including figures for "Developmental time course of social touch, parvalbumin interneurons, perineuronal nets and Mef2c expression reveals a sensitive period of somatosensory cortex development in prairie voles"

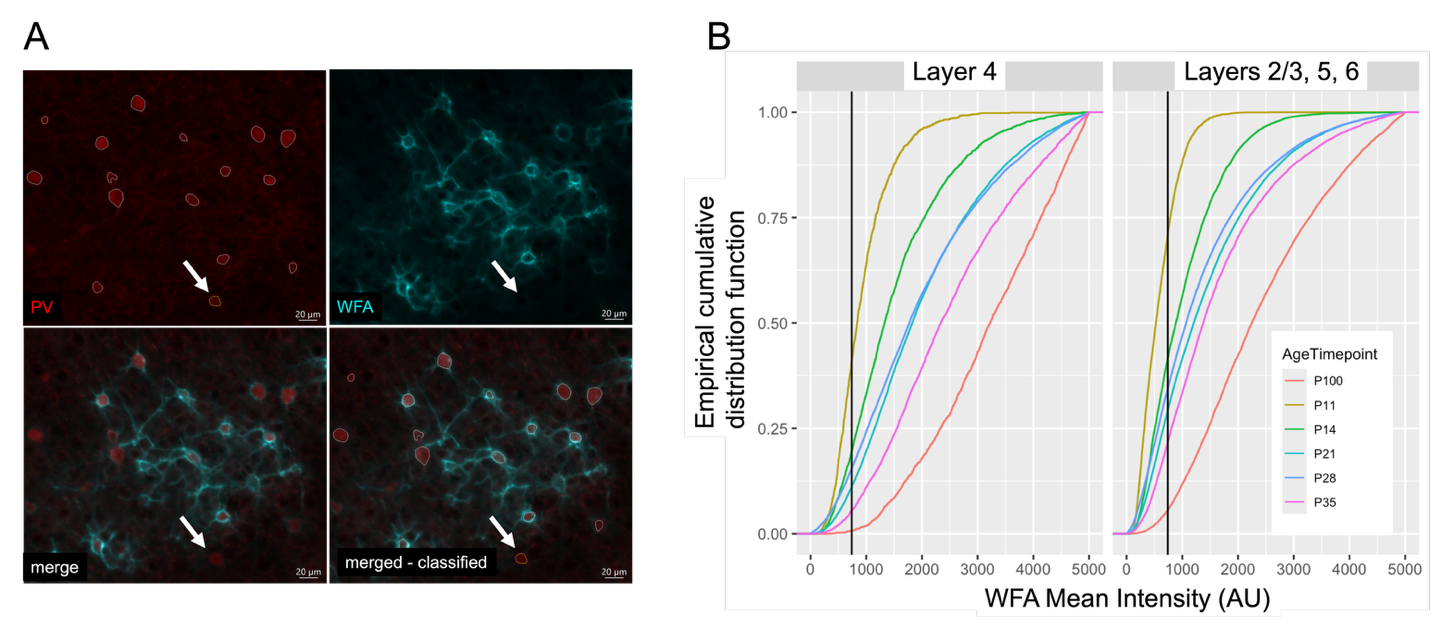

**Supplemental Figure 1 – related to Figure 2: Classification of PNNs surrounding identified PVIs**

A. Representative identified PVIs in L4 of an adult prairie vole, PV (red), WFA (cyan), merge and merge according to classification. Note one PVI (white arrow, outlined in yellow) has a mean intensity of WFA below 902 AU, and is classified as PNN-, while all other PVIs in this example are PNN+. Scale bar, 20μm.
B. Empirical cumulative distribution function (ecdf) of WFA mean intensity surrounding all PVIs according to age (color) in Layer 4 or Layer 2/3, 5, 6, black vertical line represents the 902 AU threshold for a PVI to be categorized as PNN+.

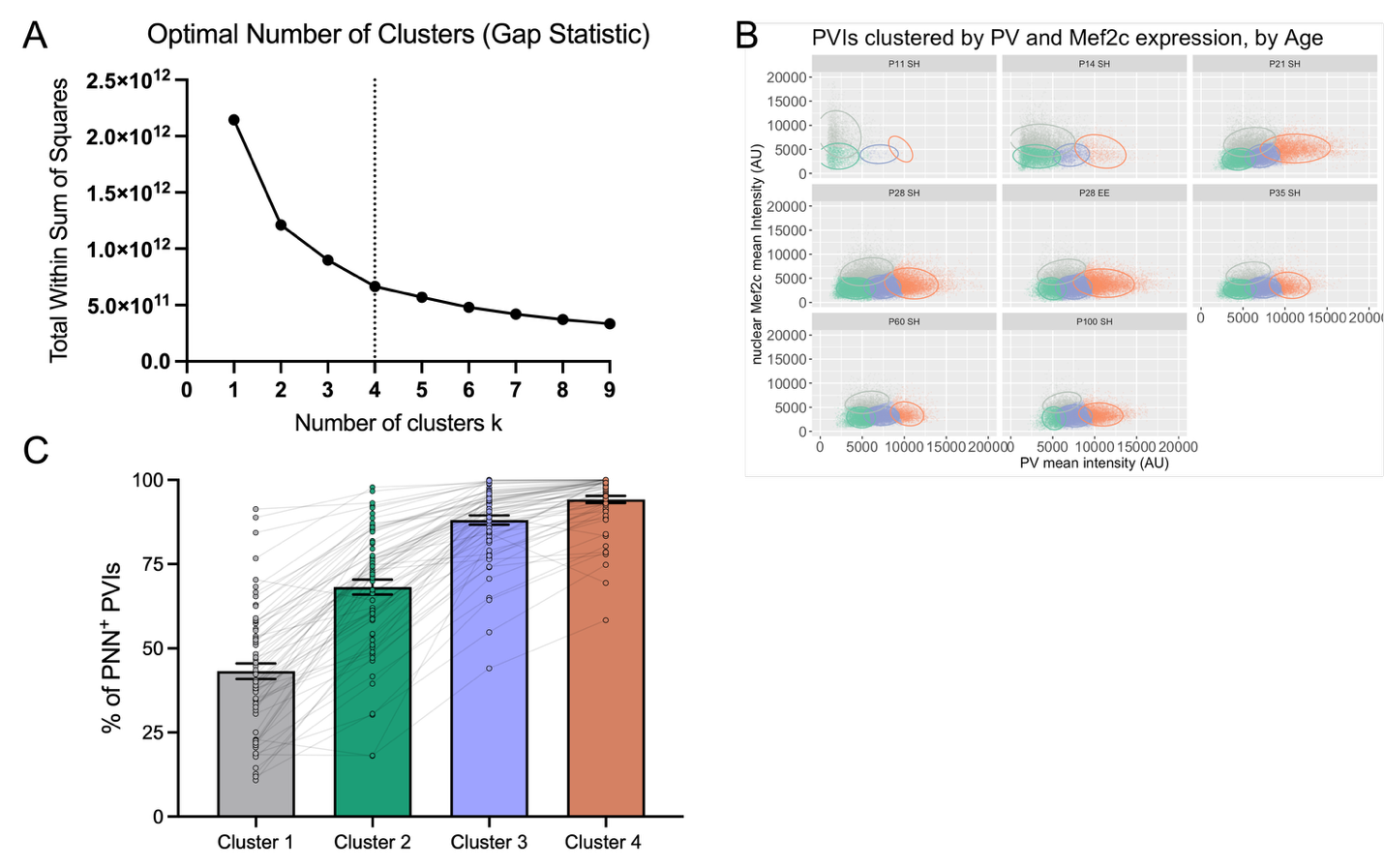

**Supplemental Figure 2 – related to Figure 4: Identification of four unique clusters of PVIs by age.**

A. To determine the optimal number of clusters k, we used the gap statistic to compare the total within cluster sum of squares (Tibshirani et al., 2001).
B. Individual PVIs clustered by PV (x-axis) and nuclear Mef2c (y-axis) expression plotted by Age timepoint and housing, standard housing (SH) or environmental enrichment (EE).
C. Proportion of PVIs with a PNN surrounding them according to cluster id. Data presented as mean +/- standard error of the mean, individual dots represent animals and lines connecting them.
